# Structural Characterization of TRAF6 N-terminal for Therapeutic Uses

**DOI:** 10.1101/2023.02.25.529911

**Authors:** Omur Guven, Halilibrahim Ciftci, Hiroshi Tateishi, Ryoko Koga, Mohamed O. Radwan, Belgin Sever, Jun-ichiro Inoue, Masami Otsuka, Mikako Fujita, Hasan DeMirci

## Abstract

Tumor Necrosis Factor Receptor Associated Factors (TRAFs) are a protein family with a wide variety of roles and binding partners. Among them, TRAF6, a ubiquitin ligase, possesses unique receptor binding specificity and shows diverse functions in immune system regulation, cellular signaling, central nervous system (CNS), and tumor formation. TRAF6 consists of an N-terminal Really Interesting New Gene (RING) domain, multiple zinc fingers, and a C-terminal TRAF domain. RING domain and zinc fingers mediate the activation of nuclear factor kappa B (NF-κB), which has essential roles in the regulation of inflammatory responses, proliferation, differentiation, migration, cell adhesion, and apoptosis. Therefore, it has been found that TRAF6 is overexpressed in various types of cancer including pancreatic, liver, lung, head and neck, breast, colorectal cancers, and melanoma along with inflammatory, autoimmune and neurodegenerative disorders. Furthermore, TRAF6 is an important therapeutic target for numerous disorders and structural studies of this protein are crucial for the development of next-generation therapeutics. Here, we present a TRAF6 N-terminal structure determined at the Turkish Light Source “*Turkish DeLight*” to 2.6 Å resolution at cryogenic temperature. This structure offers insight into the domain organization and zinc-binding, which are critical for protein function. Since the RING domain and the zinc fingers are key targets for TRAF6 therapeutics, structural insights are crucial for future research.

## 1. Introduction

Zinc finger proteins are widely distributed transcription factors in the human genome with an array of biological functions. These functions include those associated with ubiquitin-mediated protein degradation, signal transduction, differentiation, metabolism, apoptosis, autophagy, migration, invasion, and a plethora of other processes. These functions arise from the capability of zinc finger proteins to interact with their particular DNA and RNA targets. Zinc finger proteins are dependent on Zn^2+^ cations which can bind to cysteine and histidine residues. These proteins can be separated into distinct members of classical and non-classical types, referring to proteins whose zinc fingers contain the signature cys-cys-his-his (C2H2) motif and those that do not. Despite the great endeavors for the identification of the majority of zinc finger motifs, the structures of most of them have been remaining poorly characterized [1-4].

Tumor necrosis factor (TNF) receptor-associated factor (TRAF) family proteins are key regulatory molecules in the immune and inflammatory systems. They are the main signal transducers for the TNF receptor, the Interleukin-1 receptor/Toll-like receptor (IL-1/TLR), and NOD-like receptor (NLR) superfamilies. Until recently, TRAFs were classified as classical members (TRAF1-6) and a single nonclassical member (TRAF7) [5-7]. In general, TRAFs (TRAF1-7) adopt a common structure including the Really Interesting New Gene (RING) domain (except for TRAF1), zinc finger motifs, a coiled-coil domain, and a highly conserved C-terminal *β*-sandwich domain (TRAF-C or MATH domain). Apart from TRAF7, other TRAF proteins share a common structure at the TRAF-C domain. The RING domain is responsible for E3 ubiquitin ligase activity structurally supported by the zinc fingers. TRAF6 is a well-characterized E3 ubiquitin ligase, whereas the E3 activity of other TRAFs (TRAF2, TRAF3, and TRAF5) is not fully identified [8-10].

TRAF6, a non-conventional E3 ubiquitin ligase, attracts significant attention as the most studied TRAF member, differing from other TRAFs participating in the signal transduction of both the TNF receptor and the IL-1/TLR family proteins. TRAF6 is capable of activating a cascade of signaling events and upstream kinases, including kappa-B kinase (IKK), c-Jun NH_2_-terminal kinase (JNK) and p38 mitogen-activated protein kinase (MAPK) resulting in the stimulation of transcription factors including interferon-regulatory factor (IRF), the nuclear factor kappa B (NF-κB) and activator protein-1 (AP1) families [11-15]. In particular, NF-κB regulates the expression of a variety of genes involved in inflammatory responses, proliferation, differentiation, migration, cell adhesion, and apoptosis. Thus, NF-κB dysfunction can increase cancer-cell proliferation and hamper apoptosis [16-18]. Moreover, TRAF6 has been reported to enhance the ubiquitination and activation of protein kinase B (AKT) and transforming growth factor activated kinase 1 (TAK1), leading to cell cycle progression, proliferation, and migration of cancer cells along with impairment of apoptosis in cancer cells. Therefore, the overexpression of TRAF6 is linked with inflammatory disorders and various types of cancers including pancreatic, liver, lung, head and neck, breast, colorectal, prostate, melanoma, and osteosarcoma [19-24]. On the other hand, it has been also reported that TRAF6 serves important roles in osteoclastogenesis, defective lymph node organogenesis, and hypohidrotic ectodermal dysplasia [15, 25]. Recent studies also documented that high levels of TRAF6 were observed in serum patients with autoimmune diseases including systemic lupus erythematosus, rheumatoid arthritis and myasthenia gravis [26]. Besides, the connection between upregulation and/or accumulation of TRAF6 and neurodegenerative disorders such as Alzheimer’s disease, Parkinson’s disease and Amyotrophic lateral sclerosis (ALS) have been reported since TRAF6 triggers neuronal apoptosis and central nervous system (CNS) disruption. In CNS, TRAF6 also contributes to inflammatory responses in stroke and neuropathic pain [27-30].

In this study, we investigated the N-terminal region of TRAF6 at atomic resolution to examine the structure. Our findings have important implications for further pharmaceutical studies in particular for the development of next generation TRAF6 inhibitors to be effective as anti-inflammatory, anticancer, anti-osteoporosis, immunosuppressant or anti-neurodegenerative agents.

## 2. Materials and Methods

### 2.1. Transformation and expression

TRAF6 N-Terminal RING domain and 3 Zinc fingers with the sequence “MAHHHHHHHHHHVGTENLYFQSMEEIQGYDVEFDPPLESKYECPICLMALREAVQTPCGHRFCKACIIKSIRDAGHKCPVDNEILLENQLFPDNFAKREILSLMVKCPNEGCLHKMELRHLEDHQAHCEFALMDCPQCQRPFQKFHINIHILKDCPRRQVSCDNCAASMAFEDKEIHDQNCPLA” was cloned into pRSF vector with an N-terminal decahistidine purification tag with a TEV cut site. As a cloning restriction enzyme cut sites, *Hin*dIII and *Kpn*I were chosen and the Kanamycin resistance gene was used as a selection marker. The constructed plasmid was transformed into competent *Escherichia coli (E. coli)* BL21 Rosetta-2 strain, with heat shock method. Transformed bacterial cells were grown in 18 L regular LB media containing 50 µg/ml kanamycin and 35 µl/ml chloramphenicol at 37 °C. At OD600 value of 0.8, the protein expression was induced by using β-D-1-thiogalactopyranoside (IPTG) at a final concentration of 0.4 mM for 18 hours at 18 °C. Cell harvesting was done using Beckman Allegra 15R desktop centrifuge at 4 °C at 3500 rpm for 45 minutes. Cell pellets were stored at -45°C until protein purification.

### 2.2 Purification

The cells were dissolved in lysis buffer containing 500 mM NaCl, 50 mM Tris-HCl pH 8, 10% (v/v) Glycerol, 0.1% (v/v) Triton X-100, 2 mM BME and 10 µM ZnCl_2_. The homogenized cells were lysed using a Branson W250 sonifier (Brookfield, CT, USA). The cell lysate was centrifuged at 4 °C at 35000 rpm for 1 hour with Beckman Optima™ L-80XP Ultracentrifuge equipped with Ti45 rotor (Beckman, USA). The pellet containing membranes and cell debris was discarded. The supernatant containing the soluble protein was filtered through 0.2 micron hydrophilic membrane and loaded to a Ni-NTA column that was previously equilibrated with a wash buffer containing 150 mM NaCl, 20 mM Tris-HCl pH 8, 20 mM Imidazole, 10% (v/v) Glycerol, 2 mM BME and 10 µM ZnCl_2_. Unbound proteins were discarded by washing the column using a wash buffer. Then, the target protein (TRAF6 RING domain) was eluted using an elution buffer containing 250 mM NaCl, 20 mM Tris-HCl pH 8, 250 mM Imidazole, 10% (v/v) Glycerol, 2 mM BME and 10 µM ZnCl_2_. Then, the eluted TRAF6 protein was dialyzed in a dialysis membrane (3 kDa MWCO) against a buffer with the same composition as the wash buffer for 3 hours at 4 °C to remove excess imidazole. Dialyzed TRAF6 protein was cut using Tobacco Etch Virus nuclear inclusion-a endopeptidase (TEV) protease to remove the hexahistidine-tag overnight at 4 °C.

### 2.3 Crystallization

The crystallization screening of N-terminal decahistidine cleaved TRAF6 was performed using the sitting-drop microbatch under oil method against ∼3000 commercially available sparse matrix crystallization screening conditions in a 1:1 volumetric ratio in 72-Terasaki plates (Greiner Bio-One, Kremsmünster, Austria) as described in Ertem 2021 et al. [31]. The mixtures were covered with 16.6 µl 100% paraffin oil (Tekkim Kimya, Istanbul, Türkiye). The crystallization plates were incubated at 4 °C and checked frequently under a stereo light microscope. The best TRAF6 crystals were grown within one month in Salt Rx-I condition #22 (Hampton Research, USA). This condition contains 1.2 M sodium citrate tribasic dihydrate and 0.1 M TRIS-HCl 8.5.

### 2.4 Crystal harvesting and delivery

The TRAF6 crystals were harvested using MiTeGen MicroLoops attached to a magnetic wand [32] while being monitored under microscope [33]. The obtained crystals were flash frozen by plunging in liquid nitrogen and placed in a cryo-cooled sample storage puck (Cat#M-CP-111-021, MiTeGen, USA). Then, the puck was placed into the *Turkish DeLight* liquid nitrogen-filled autosample dewar at 100 °K.

### 2.5 Data collection and data reduction

Collection of diffraction data from the TRAF6 crystal was performed by utilizing Rigaku’s XtaLAB Synergy Flow XRD source “*Turkish DeLight*” at University of Health Sciences (Istanbul, Türkiye) with *CrysAlisPro* software 1.171.42.35a [34]. The crystals were kept cooled by the Cryostream 800 Plus system, which was set to 100 °K. The PhotonJet-R X-ray generator operated at 30 mA, 1200.0 W and 40 kV with 10% beam intensity. The data was collected at 1.54Å wavelength and the detector distance was set to 47.00 mm. The crystal oscillation width was set to 0.25 degrees per image while exposure time was 20.0 minutes. CrysAlis Pro version 1.171.42.35a [34] was utilized to perform data reduction and an *.mtz file was obtained as the result.

### 2.6 Structure determination and refinement

The cryogenic TRAF6 structure was determined at 2.6 Å with the space group P1 utilizing *PHASER* 2.8.3 [35], an automated molecular replacement program within the *PHENIX* suite 1.20.1 [36]. The previously released TRAF6 structure with PDB ID: 3HCS was used as an initial search model for molecular replacement [37]. Simulated annealing and rigid-body refinements were performed during the first refinement cycle including individual coordinates and translation/liberates/screw (TLS) parameters were refined. The structure was checked by *COOT* [38] after each set of refinement and the solvent molecules were added into unfilled, appropriate electron density maps. The obtained final structure was examined by using *PyMOL* [39] 2.5.4 and *COOT* 0.9.8, and the figures were created. Data collection and refinement statistics were given in Table 1.

**Table 1.**
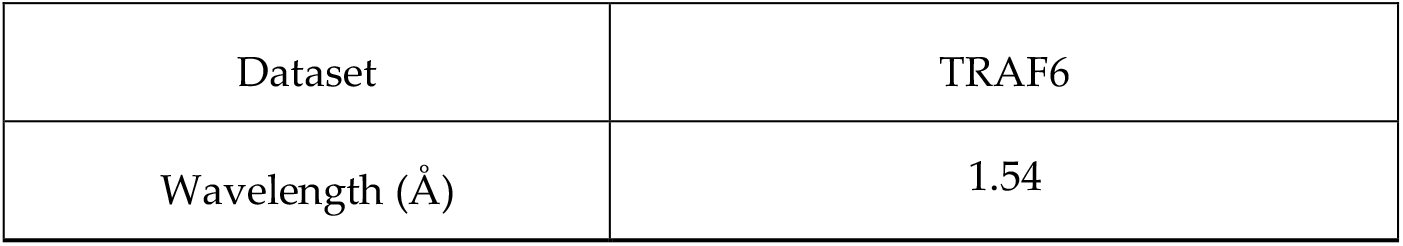
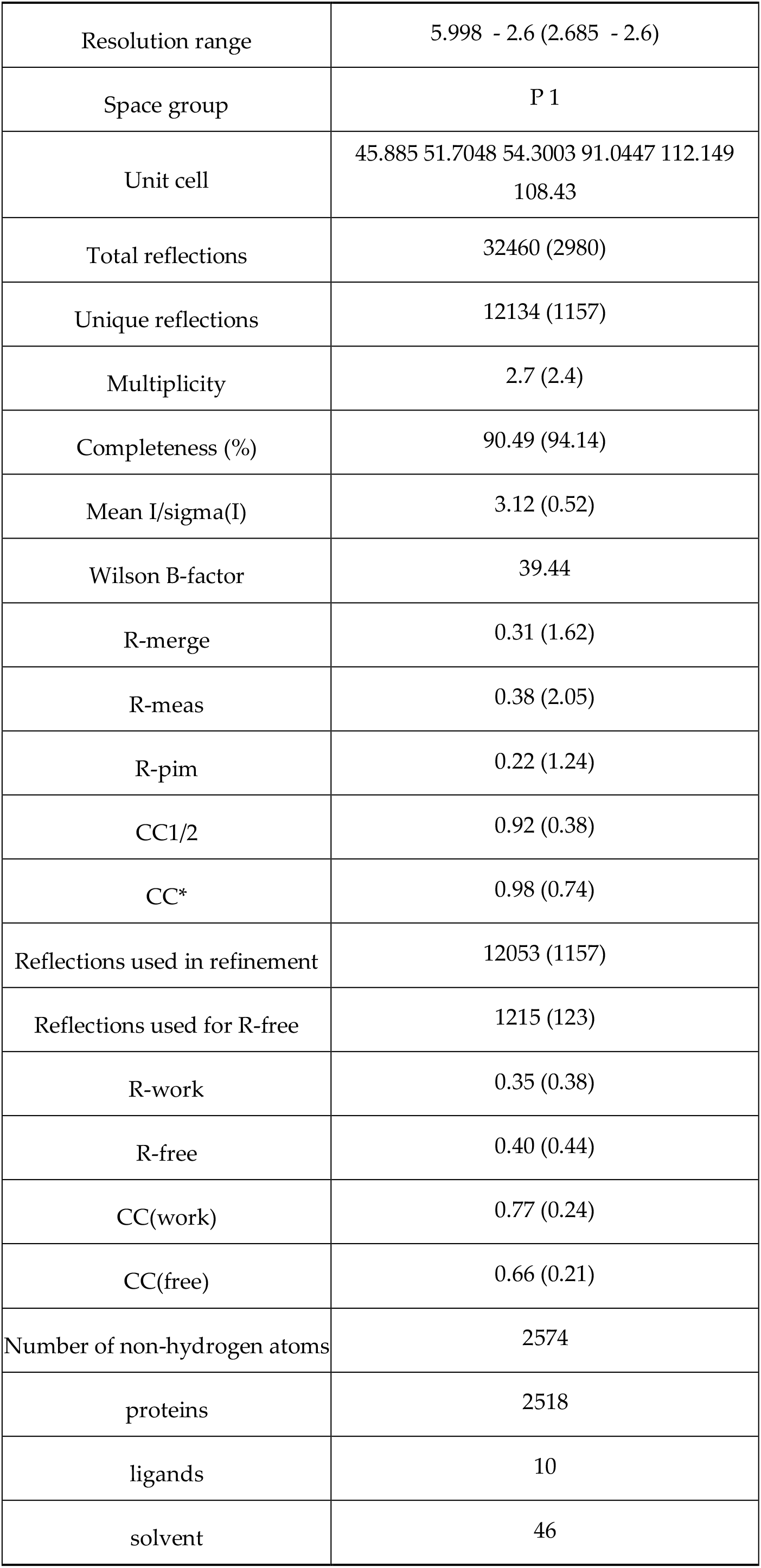
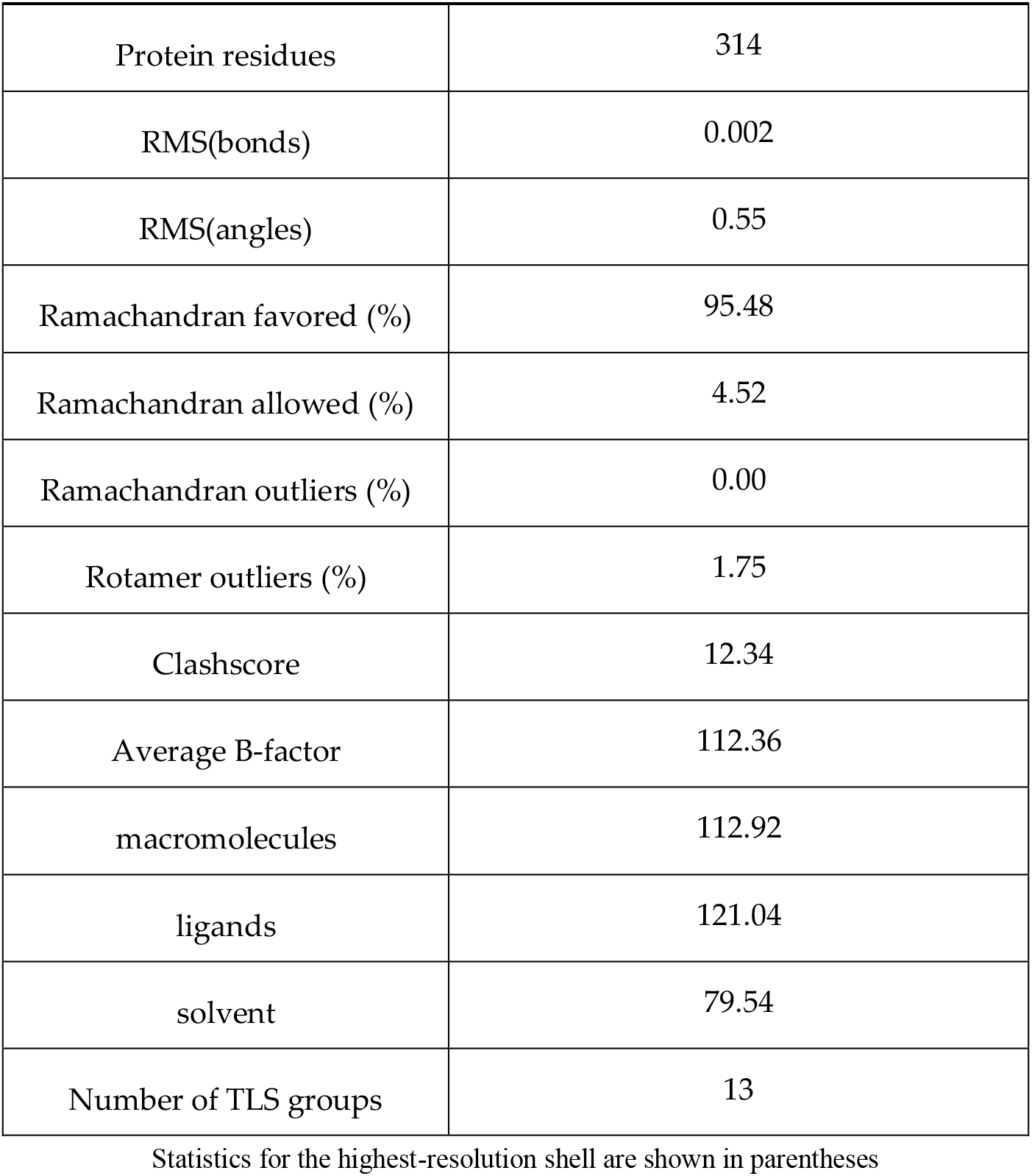
Data collection and refinement statistics

| Dataset | TRAF6 |
| --- | --- |
| Wavelength (Å) | 1.54 |
| Resolution range | 5.998 - 2.6 (2.685 - 2.6) |
| Space group | P 1 |
| Unit cell | 45.885 51.7048 54.3003 91.0447 112.149<br>108.43 |
| Total reflections | 32460 (2980) |
| Unique reflections | 12134 (1157) |
| Multiplicity | 2.7 (2.4) |
| Completeness (%) | 90.49 (94.14) |
| Mean I/sigma(I) | 3.12 (0.52) |
| Wilson B-factor | 39.44 |
| R-merge | 0.31 (1.62) |
| R-meas | 0.38 (2.05) |
| R-pim | 0.22 (1.24) |
| CC1/2 | 0.92 (0.38) |
| CC* | 0.98 (0.74) |
| Reflections used in refinement | 12053 (1157) |
| Reflections used for R-free | 1215 (123) |
| R-work | 0.35 (0.38) |
| R-free | 0.40 (0.44) |
| CC(work) | 0.77 (0.24) |
| CC(free) | 0.66 (0.21) |
| Number of non-hydrogen atoms | 2574 |
| proteins | 2518 |
| ligands | 10 |
| solvent | 46 |
| Protein residues | 314 |
| RMS(bonds) | 0.002 |
| RMS(angles) | 0.55 |
| Ramachandran favored (%) | 95.48 |
| Ramachandran allowed (%) | 4.52 |
| Ramachandran outliers (%) | 0.00 |
| Rotamer outliers (%) | 1.75 |
| Clashscore | 12.34 |
| Average B-factor | 112.36 |
| macromolecules | 112.92 |
| ligands | 121.04 |
| solvent | 79.54 |
| Number of TLS groups | 13 |

Statistics for the highest-resolution shell are shown in parentheses

## 3. Results

### 3.1. TRAF6 N-terminal structure was determined at 2.6 Å resolution at Turkish DeLight

TRAF6 is a 59.4 kDa protein consisting of 522 amino acids. Here we structurally calculated the 18.14 kDa N-terminal region consisting of 157 amino acids. The crystal belongs to the P1 triclinic space group with a=45.885, b=51.7048, c=54.3003, α=91.04, β=112.15, γ=108.43. The dimerized structure of TRAF6 N-terminal region was determined to 2.6 Å resolution at cryogenic temperature at the Turkish Light Source “*Turkish DeLight*” [40] (Figure 1). The determined structure was deposited to PDB database with the ID: 8HZ2.

**Figure 1.**
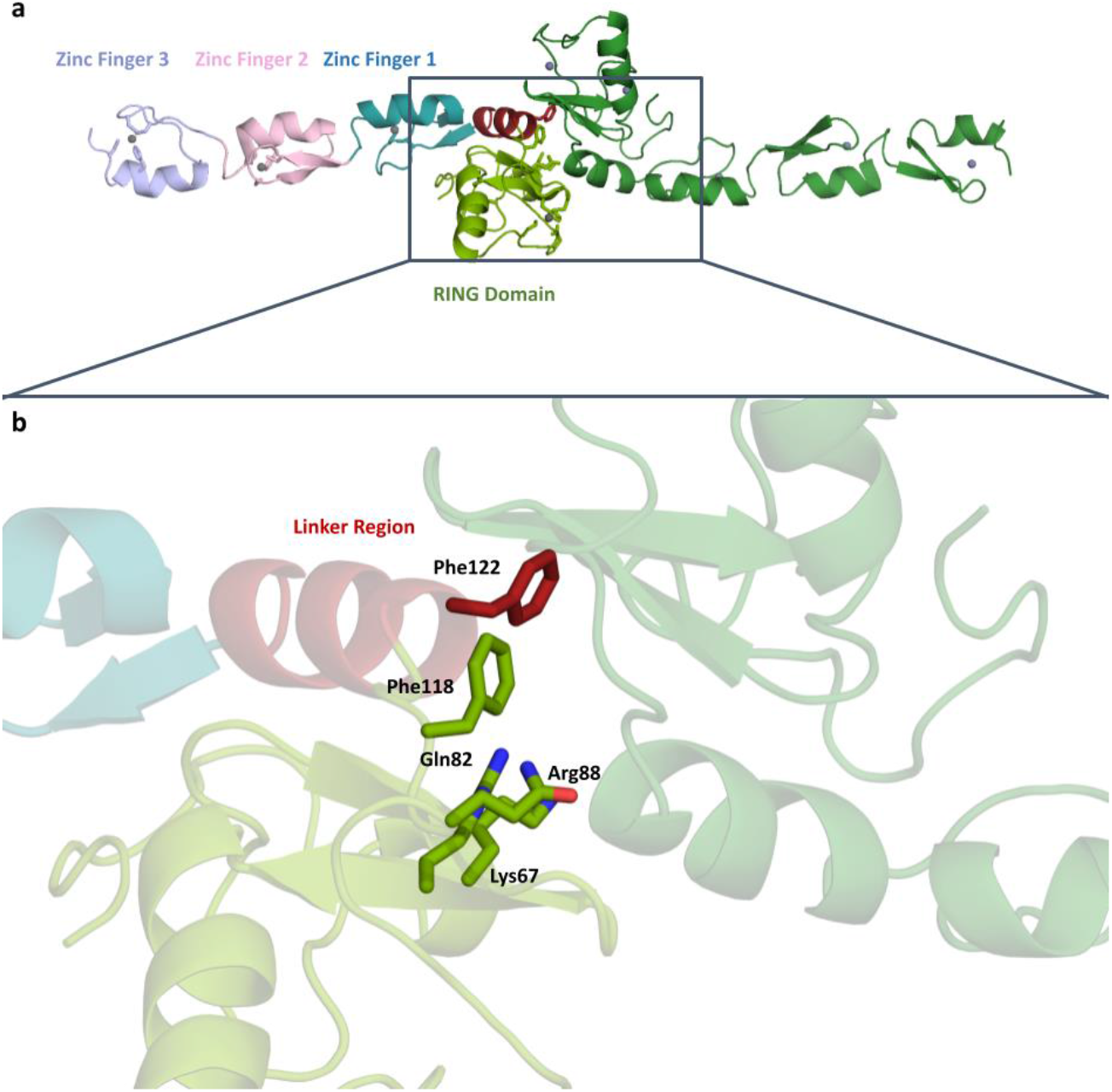
**(a)** Dimeric structure of TRAF6 N-terminal region at 2.6 Å (Chain A RING Domain: Splitpea, ZF1: Deepteal, ZF2: Lightpink, ZF3: Lightblue, Zn: Gray ; Chain B: Forest, Zinc atoms: Gray) and domain organization of TRAF6 N-terminal region. **(b)** A close-up of the dimerization residues are shown at the dimerization surface.

Our study reveals the structure of TRAF6, which has dimerized at the N-terminal RING domain and linker region. Residues known to mediate the dimerization are shown in Figure 1. There are four residues in the RING domain (Lys67, Gln82, Arg88 and Phe118) and one residue in the linker region (Phe122) shown participating in dimerization.

TRAF6 N-terminal region consists of 5 domains, a RING domain, a linker helix, and three zinc fingers. Figure 1a shows the domain organization in detail, the zinc atoms can be clearly seen at the center of the domains.

### 3.2. Detailed analysis of RING domain and zinc fingers

A closer look at the RING domain (Figure 2) and zinc fingers (Figure 3) was also taken. Every zinc atom and their interacting residues are visible within the refined *2Fo-Fc* density map. A clear repeating pattern among the zinc fingers can be observed in secondary structures. Each finger is made of a Sheet-Loop-Sheet-Helix-Loop pattern. Besides, there are always three Cys residues and one His residues forming the finger. While the two cysteine residues are located on the first loop, the third cysteine is located on the second loop and the histidine is located on the helix region. The distances between zinc and interacting residues are also conserved, ranging from 2.0 to 2.3 Å. We have generated the *2Fo-Fc* electron density maps of the zinc and the interacting residues which show continuation throughout and enclosing the zinc atom and the residues.

**Figure 2.**
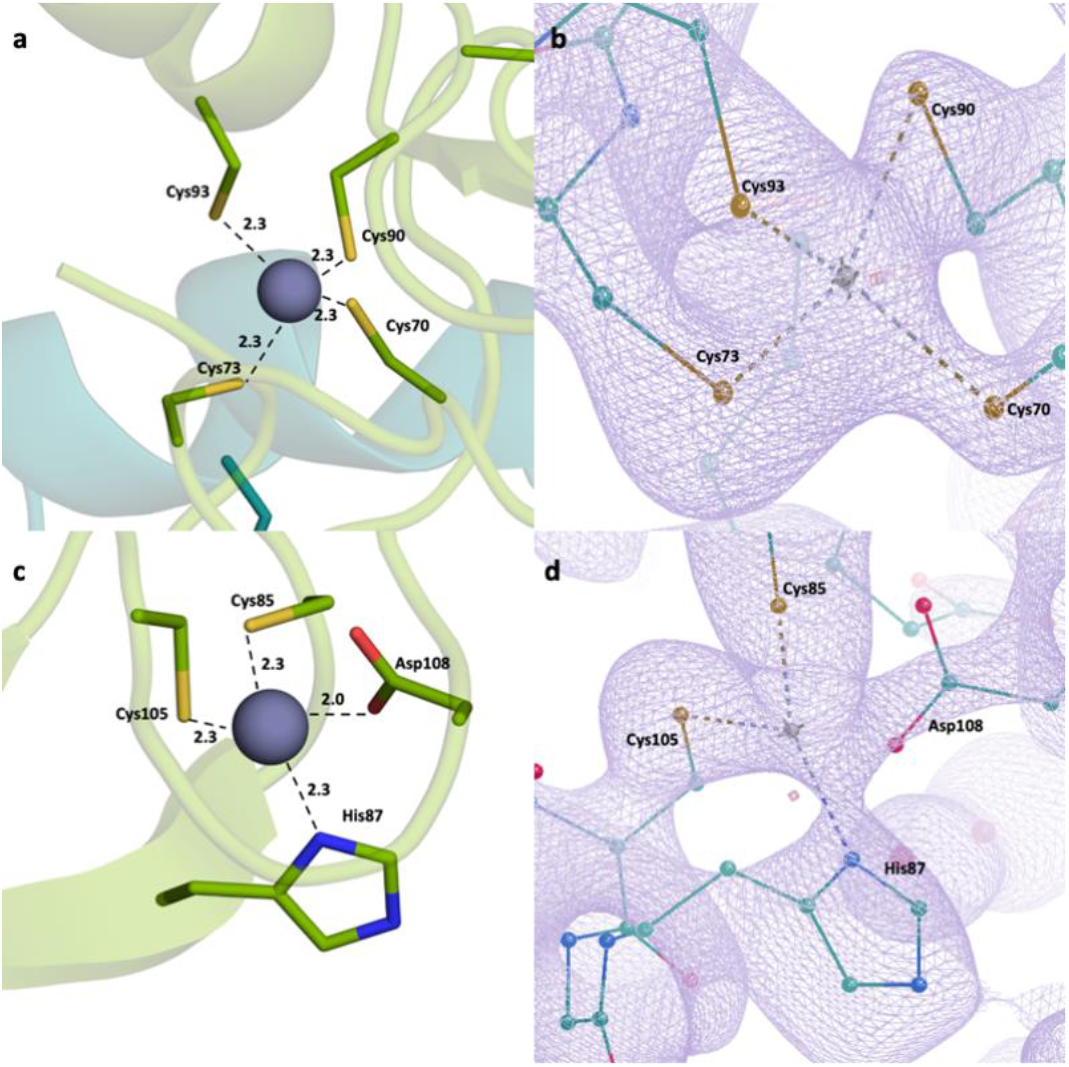
RING domain zinc-interacting residues. **(a), (c)** Distances between residues and the bivalent zinc ions.**(b), (d)** The 2Fo-Fc electron density maps shown at sigma level of 1 RMSD showing the interaction.

**Figure 3.**
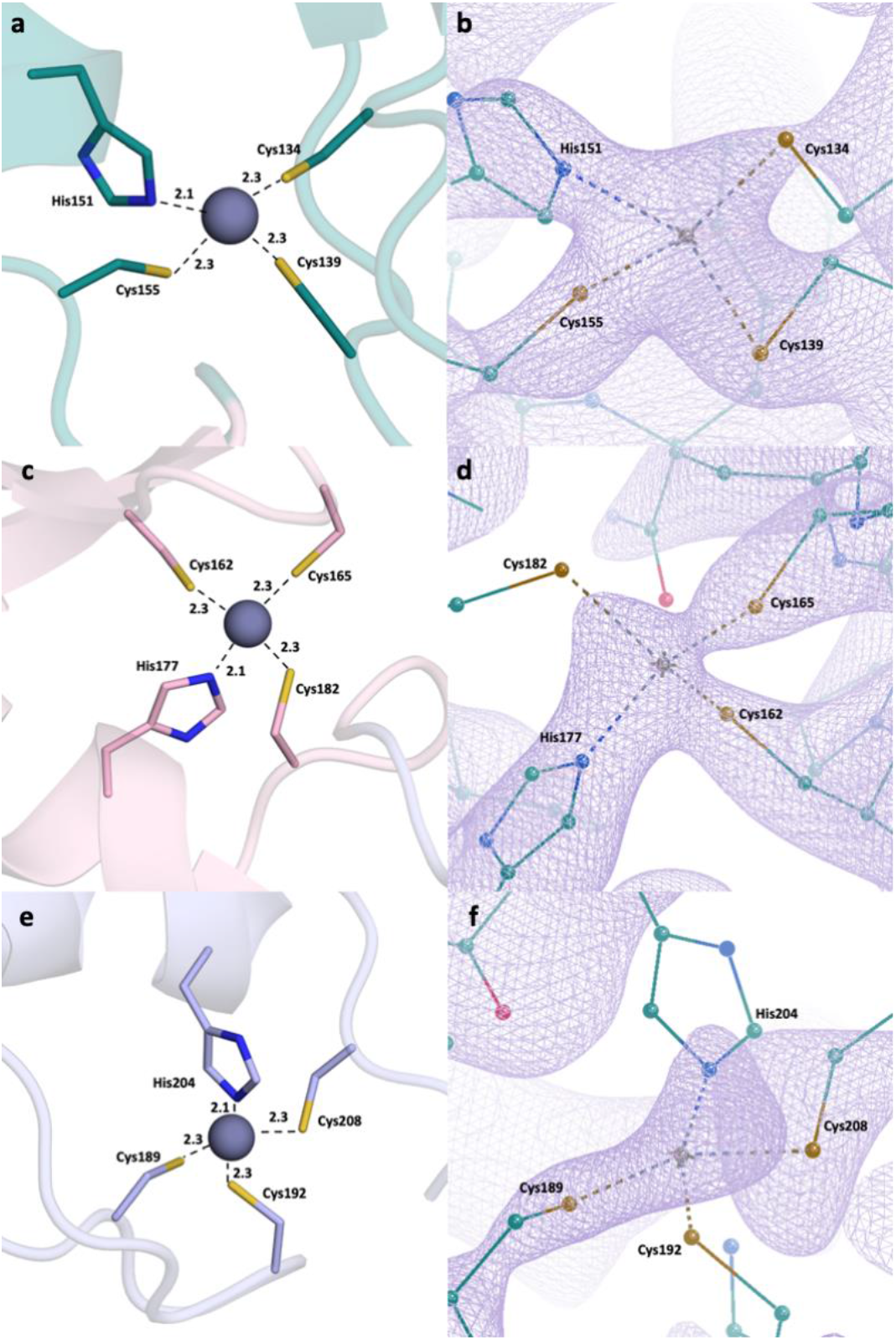
Zinc fingers and zinc-interacting residues. **(a)** Zinc finger 1 distances. **(b)** Zinc finger 1 electron density map at sigma level of 1 RMSD. **(c)** Zinc finger 2 distances. **(d)** Zinc finger 2 *2Fo-Fc* electron density map at sigma level of 1 RMSD. **(e)** Zinc finger 3 distances. **(f)** Zinc finger 3 *2Fo-Fc* electron density map at sigma level of 1 RMSD.

**Figure 4.**
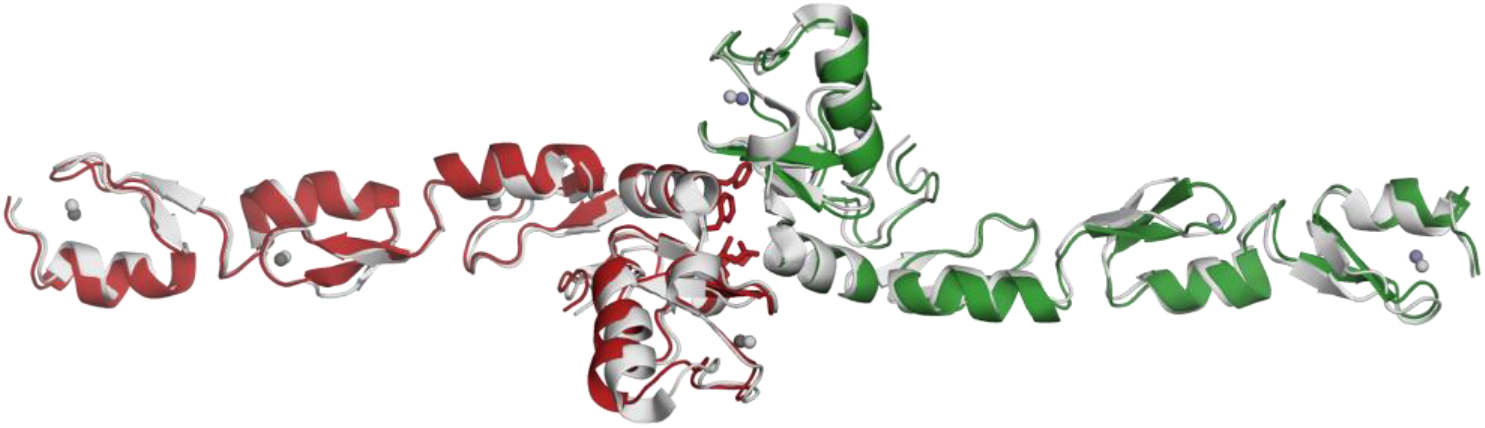
Structure alignment between the obtained data (PDB ID: 8HZ2) and the used model (PDB ID: 3HCS). Model structure is shown in gray.

### 3.3 Structural alignment with the reference protein

Our structure (PDB ID:8HZ2) and the reference structure (PDB ID:3HCS) are aligned with the RMSD value of 1.09, and show minor differences within the loops. Also, we have looked at the zinc-interacting residues in both structures. (Supplementary Figure 1) Although the residues have similar conformations, there are slight differences based on the loop movements. We have also checked the distances between the zinc-binding residues and zinc ions. Here we see that these distances are similar in reference structure and our structure, although the former have a slightly larger range, between 2.1 and 2.8Å. (data not shown).

## 4. Discussion

TRAF6 is widely distributed in the brain, lung, liver, skeletal muscle and kidney and is involved in a great number of immune and inflammatory reactions as a characteristic E3 ubiquitin ligase. TRAF6 plays a pivotal role in NF-κB stimulation, which triggers a vast array of cellular and organismal processes such as development, immunity, tissue homeostasis and inflammation regulating gene expression, apoptosis and proliferation at molecular and cellular levels. Therefore, TRAF6 is associated with diverse abnormalities including different cancer types, autoimmune diseases, neurodegenerative disorders and inflammatory diseases [41-43].

Having access to the structural details of TRAF6 provides insights that can support the development of new generation anticancer therapeutics. Furthermore, the previously deposited structure used as a model (PDB ID: 3HCS) serves as a reference point from which we can observe that our structure is highly similar. The most significant characteristic of TRAF6 is the zinc interaction. Bivalent ions provide a structural scaffold around where the zinc fingers and the RING domain folds. Therefore, zinc has an important role in TRAF6 function, and altering the zinc interaction is a good starting point for inhibition. The overall structure of the TRAF6 N-terminal region shows the dimerized structure with zinc atoms at the center of each domain. The domain organization shows, in addition to RING domain and zinc fingers, a linker region, consisting of a single helix (Figure 3). This region has a role in N-terminal dimerization, as a phenylalanine residue here is part of the dimerization surface.

RING domain and zinc fingers were analyzed in detail. Zinc fingers are one of the most abundant structural motifs observed in proteins [44]. As their name suggests, they are characterized around a bivalent zinc ion and they can interact with a wide range of molecules, such as nucleic acids and other proteins. Therefore, zinc finger proteins have a wide variety of functions, from transcriptional regulation to actin targeting. Zinc fingers have common patterns, for example, they consist mainly of cysteine and histidine residues, in different ratios [45]. Classical zinc fingers have two of each (Cys_2_His_2_), however, some cases show differences. Zinc finger domains have two β-sheets and one ɑ-helix, although the number of loops change depending on the protein.

RING domain is a common domain in ubiquitin-ligase proteins (E3), with over 340 such proteins possessing this domain. It interacts with two bivalent zinc ions, forming RING fingers [46]. RING domains interact with DNA, therefore, the proteins including RING domain could mediate DNA transcription. The presented structure has a RING domain and three zinc fingers. Each zinc finger follows a conserved pattern; Sheet-Loop-Sheet-Helix-Loop. Moreover, each zinc finger has the same residue pattern, two cysteine residues in the first loop, one histidine residue in the helix and a third cysteine residue in the second loop. This is an expected result, as the zinc fingers fold around bivalent zinc ions, and they possess similar patterns. This is a classical zinc finger pattern in a way that it has two sheets and a helix, however, it differs as classical zinc finger in its Cys_2_His_2_ structure.

We have also investigated the distances between the zinc-interacting-residues and the zinc ion. We observed that the distances are well within the range of strong bond formation at up to 2.4 Å. The cysteine residues interact with the zinc through the sulfur at the side chain while the histidines interact through their side chain nitrogens. We compared the distances in three zinc fingers, and we observed that they are similar as well, further showing that the zinc fingers have common characteristics. Moreover, it is confirmed that our structure has a RING finger, with its two bivalent zinc ions present thus, forming a RING domain. This domain mediates DNA interaction, and is crucial for the role of TRAF6 on NF-κB regulation. These four domains, all revolving around the central zinc atom, are great targets for TRAF6 therapeutics. The high number of cysteine residues result in disulfide bond formation, in addition to zinc interaction, therefore altering these bonds with reducing agents (for example, dithiol compounds) will go a long way in modulating TRAF6 function.

Structural alignment with the model protein was performed. We have used a deposited structure (PDB ID: 3HCS) as a model, and have aligned it to the obtained structure (Figure 6). We observed that the structures aligned with high similarity (RMSD= 1.09), except for the loops. This is expected as loops are generally flexible, and the difference is based on the natural properties of the protein. We have also looked into the RING domain and zinc fingers in detail. We observed that the distances between zinc-interacting residues and zinc ions are very similar to that of obtained structure (PDB ID: 8HZ2), ranging between 2.1 and 2.8Å. This is an expected result as the zinc fingers are conserved regions further confirmed by these distances.

Over the years, our research group has also pursued the discovery of small molecules with efficacy on zinc finger proteins. With great effort in this area, we have discovered compounds with dimethylaminopyridine and histidine nuclei and proved their remarkable efficacy against zinc finger proteins such as human immunodeficiency virus type I enhancer binding protein 1 (HIV-EP1) [47,48]. Following these efforts, SN-1 (Figure 7), developed by us as an inhibitor of zinc finger transcription factor [49], was found to enhance steady-state expression level of antiviral apolipoprotein B mRNA-editing enzyme-catalytic polypeptide-like (APOBEC) 3G (A3G) bearing two zinc-binding domains in the presence of viral infectivity factor (Vif) protein [50]. Then, we also reported that SN-1 bound to TRAF6 suppressing its auto-ubiquitination and downstream NF-κB signaling supported by the molecular docking studies exhibiting that SN-1 interacted with the first zinc finger of TRAF6 [51]. Besides, we further demonstrated that SN-1 derivatives hold promise for developing new drug candidates targeting zinc proteins [54]. In future studies, we plan to perform structural studies of TRAF6 with SN-1.

**Figure 7.**
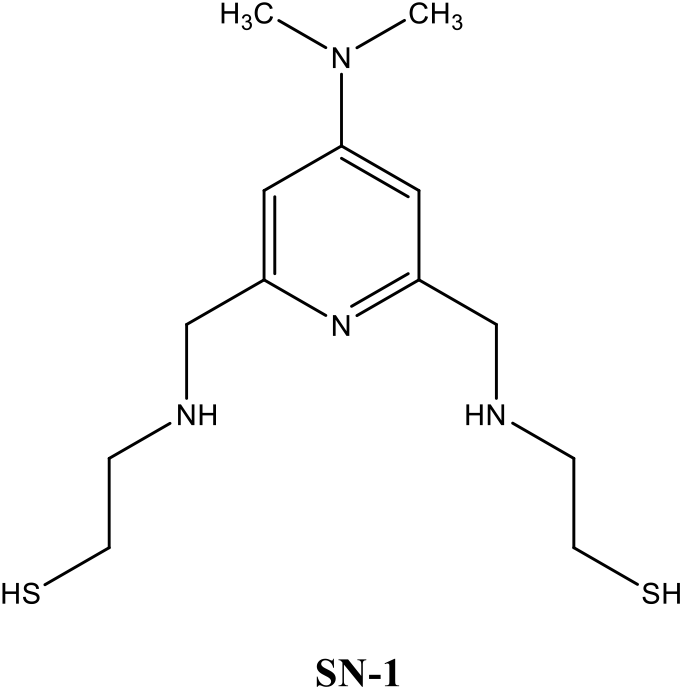
The chemical structure of SN-1

## 5. Conclusions

TRAF6 serves important roles in a variety of physiological and pathological processes resulting in cancer, inflammation, neurodegeneration, osteoporosis, autoimmunity and many other disorders. Therefore, examining and knowing the accurate protein structure can guide us in understanding the exact mechanism of action. The current research manifested the characteristics of the RING domain organization and zinc-binding of TRAF6, shedding light on the crucial functions of the protein. This study will also encourage further evaluation of the structure of TRAF6 with our previously developed compound SN-1, a potential TRAF6 inhibitor. SN-1 could be a lead compound for the enhancement of the specificity to TRAF6 for future studies. Based on our continuous endeavors, we aim to develop TRAF6-specific drug candidates to be effective against various disorders.

## Supporting information

https://drive.google.com/drive/u/0/folders/1GejcUMXvZ9ryu4zG1ClzIVw0CU0_s54A

## Supplementary Materials

The following supporting information can be downloaded at: www.mdpi.com/xxx/s1, Figure S1: A close up of zinc-binding pockets in aligned structure

## Author Contributions

Conceptualization, Ö.G., H.C., R.K. and H.D.; methodology, Ö.G., H.C. and H.D.; software, Ö..G, H.C. and H.D.; validation, Ö.G., H.C. and H.D.; formal analysis, Ö.G., H.C. and H.D.; investigation, Ö.G., H.C., J.I. and H.D.; resources, Ö.G., H.C. and H.D.; data curation, Ö.G., H.C. and H.D; writing—original draft preparation, Ö.G, H.C. and H.D; writing—review and editing, Ö.G., H.C., M.O.R., B.S., H.T., M.O., M.F. and H.D.; visualization, Ö.G., H.C. and H.D.; supervision, H.D.; project administration, H.D.; funding acquisition, H.C. and H.D. All authors have read and agreed to the published version of the manuscript.

## Data Availability Statement

The Traf6 N-terminal presented in this manuscript has been deposited to the Protein Data Bank under the accession number 8HZ2 Any remaining information can be obtained from the corresponding author upon request.

## Acknowledgements

Authors would like to dedicate this manuscript to the memory of Dr. Albert E. Dahlberg and Dr. Nizar Turker. The authors gratefully acknowledge use of the services and Turkish Light Source (Turkish DeLight) X-ray facility at Sağlık Bilimleri University Deneysel Tıp Araştırma ve Uygulama Merkezi (SBU-DETUAM).

## References

1. Laity, J.H.; Lee, B.M.; Wright, P.E. Zinc finger proteins: new insights into structural and functional diversity. Curr Opin Struct Biol 2001, 11, 39–46, doi: 10.1016/s0959-440x(00)00167-6.

2. Jen, J.; Wang, Y.C. Zinc finger proteins in cancer progression. J Biomed Sci 2016, 23, 53, doi: 10.1186/s12929-016-0269-9.

3. Cassandri, M.; Smirnov, A.; Novelli, F.; Pitolli, C.; Agostini, M.; Malewicz, M.; Melino, G.; Raschellà, G. Zinc-finger proteins in health and disease. Cell Death Discov 2017, 3, 17071, doi: 10.1038/cddiscovery.2017.71.

4. Bu, S.; Lv, Y.; Liu, Y.; Qiao, S.; Wang, H. Zinc Finger Proteins in Neuro-Related Diseases Progression. Front Neurosci 2021, 15, 760567, doi: 10.3389/fnins.2021.760567.

5. Zhao, L.; Hao, Y.; Song, Z.; Fan, Y.; Li, S. TRIM37 negatively regulates inflammatory responses induced by virus infection via controlling TRAF6 ubiquitination. Biochem Biophys Res Commun 2021, 556, 87–92, doi: 10.1016/j.bbrc.2021.03.147.

6. Li, J.; Liu, N.; Tang, L.; Yan, B.; Chen, X.; Zhang, J.; Peng, C. The relationship between TRAF6 and tumors. Cancer Cell Int 2020, 20, 429, doi: 10.1186/s12935-020-01517-z.

7. Chen, Y.; Li, Y.; Li, P.T.; Luo, Z.H.; Zhang, Z.P.; Wang, Y.L.; Zou, P.F. Novel Findings in Teleost TRAF4, a Protein Acts as an Enhancer in TRIF and TRAF6 Mediated Antiviral and Inflammatory Signaling. Front Immunol 2022, 13, 944528, doi: 10.3389/fimmu.2022.944528.

8. Lamothe, B.; Campos, A.D.; Webster, W.K.; Gopinathan, A.; Hur, L.; Darnay, B.G. The RING domain and first zinc finger of TRAF6 coordinate signaling by interleukin-1, lipopolysaccharide, and RANKL. J Biol Chem 2008, 283, 24871–24880, doi: 10.1074/jbc.M802749200.

9. He, X.; Li, Y.; Li, C.; Liu, L.J.; Zhang, X.D.; Liu, Y.; Shu, H.B. USP2a negatively regulates IL-1β- and virus-induced NF-κB activation by deubiquitinating TRAF6. J Mol Cell Biol 2013, 5, 39–47, doi: 10.1093/jmcb/mjs024.

10. Lalani, A.I.; Zhu, S.; Gokhale, S.; Jin, J.; Xie, P. TRAF molecules in inflammation and inflammatory diseases. Curr Pharmacol Rep 2018, 4, 64–90, doi: 10.1007/s40495-017-0117-y.

11. Bradley, J.R.; Pober, J.S. Tumor necrosis factor receptor-associated factors (TRAFs). Oncogene 2001, 20, 6482–6491, doi: 10.1038/sj.onc.1204788.

12. Wang, P.H.; Wan, D.H.; Gu, Z.H.; Deng, X.X.; Weng, S.P.; Yu XQ, He, J.G. Litopenaeus vannamei tumor necrosis factor receptor-associated factor 6 (TRAF6) responds to Vibrio alginolyticus and white spot syndrome virus (WSSV) infection and activates antimicrobial peptide genes. Dev Comp Immunol 2011, 35, 105–114, doi: 10.1016/j.dci.2010.08.013.

13. Walsh, M.C.; Lee, J.; Choi, Y. Tumor necrosis factor receptor-associated factor 6 (TRAF6) regulation of development, function, and homeostasis of the immune system. Immunol Rev 2015, 266, 72–92, doi: 10.1111/imr.12302.

14. Inoue, J.I.; Ishida, T.; Tsukamoto, N.; Kobayashi, N.; Naito, A.; Azuma, S.; Yamamoto, T. Tumor necrosis factor receptor-associated factor (TRAF) family: adapter proteins that mediate cytokine signaling. Exp Cell Res 2000, 254, 14–24, doi: 10.1006/excr.1999.4733.

15. Yamamoto, M.; Gohda, J.; Akiyama, T.; Inoue, J.I. TNF receptor-associated factor 6 (TRAF6) plays crucial roles in multiple biological systems through polyubiquitination-mediated NF-κB activation. Proc Jpn Acad Ser B Phys Biol Sci 2021, 97, 145–160, doi: 10.2183/pjab.97.009.

16. Hayden, M.S.; Ghosh, S. Signaling to NF-kappaB. Genes Dev 2004, 18, 2195–2224, doi: 10.1101/gad.1228704.

17. Park, M.H.; Hong, J.T. Roles of NF-κB in Cancer and Inflammatory Diseases and Their Therapeutic Approaches. Cells 2016, 5, 15, doi: 10.3390/cells5020015.

18. Soleimani, A.; Rahmani, F.; Ferns, G.A.; Ryzhikov, M.; Avan, A.; Hassanian, S.M. Role of the NF-κB signaling pathway in the pathogenesis of colorectal cancer. Gene 2020, 726, 144132, doi: 10.1016/j.gene.2019.144132.

19. Middleton, A.J.; Budhidarmo, R.; Das, A.; Zhu, J.; Foglizzo, M.; Mace, P.D.; Day, C.L. The activity of TRAF RING homo- and heterodimers is regulated by zinc finger 1. Nat Commun 2017, 8, 1788, doi: 10.1038/s41467-017-01665-3.

20. Qi, Y.; Pradipta, A.R.; Li, M.; Zhao, X.; Lu, L.; Fu, X.; Wei, J.; Hsung, R.P.; Tanaka, K.; Zhou, L. Cinchonine induces apoptosis of HeLa and A549 cells through targeting TRAF6. J Exp Clin Cancer Res 2017, 36, 35, doi: 10.1186/s13046-017-0502-8.

21. Khusbu, F.Y.; Zhou, X.; Roy, M.; Chen, F.Z.; Cao, Q.; Chen, H.C. Resveratrol induces depletion of TRAF6 and suppresses prostate cancer cell proliferation and migration. Int J Biochem Cell Biol 2020, 118, 105644, doi: 10.1016/j.biocel.2019.105644. E

22. Li, N.; Luo, L.; Wei, J.; Liu, Y.; Haque, N.; Huang, H.; Qi, Y.; Huang, Z. Identification of a new TRAF6 inhibitor for the treatment of hepatocellular carcinoma. Int J Biol Macromol 2021, 182, 910–920, doi: 10.1016/j.ijbiomac.2021.04.081.

23. Guangwei, Z.; Zhibin, C.; Qin, W.; Chunlin, L.; Penghang, L.; Ruofan, H.; Hui, C.; Hoffman, R.M.; Jianxin, Y. TRAF6 regulates the signaling pathway influencing colorectal cancer function through ubiquitination mechanisms. Cancer Sci 2022, 113, 1393–1405, doi: 10.1111/cas.15302.

24. Zhao, X.; Ren, L.; Wang, X.; Han, G.; Wang, S.; Yao, Q.; Qi, Y. Benzoyl-xanthone derivative induces apoptosis in MCF-7 cells by binding TRAF6. Exp Ther Med 2022, 23, 181, doi: 10.3892/etm.2021.11104.

25. Bai, S.; Zha, J.; Zhao, H.; Ross, F.P.; Teitelbaum, S.L. Tumor necrosis factor receptor-associated factor 6 is an intranuclear transcriptional coactivator in osteoclasts. J Biol Chem 2008, 283, 30861–30867, doi: 10.1074/jbc.M802525200.

26. Li, T.; Li, Y.; Li, J.W.; Qin, Y.H.; Zhai, H.; Feng, B.; Li, H.; Zhang, N.N.; Yang, C.S. Expression of TRAF6 in peripheral blood B cells of patients with myasthenia gravis. BMC Neurol 2022, 22, 302, doi: 10.1186/s12883-022-02833-9.

27. Semmler, S.; Gagné, M.; Garg, P.; Pickles, S.R.; Baudouin, C.; Hamon-Keromen, E.; Destroismaisons, L.; Khalfallah, Y.; Chaineau, M.; Caron, E.; Bayne, A.N.; Trempe, J.F.; Cashman, N.R.; Star, A.T.; Haqqani, A.S.; Durcan, T.M.; Meiering, E.M.; Robertson, J.; Grandvaux, N.; Plotkin, S.S.; McBride, H.M.; Vande Velde, C. TNF receptor-associated factor 6 interacts with ALS-linked misfolded superoxide dismutase 1 and promotes aggregation. J Biol Chem 2020, 295, 3808–3825, doi: 10.1074/jbc.RA119.011215.

28. Huang, H.; Xia, A.; Sun, L.; Lu, C.; Liu, Y.; Zhu, Z.; Wang, S.; Cai, J.; Zhou, X.; Liu, S. Pathogenic Functions of Tumor Necrosis Factor Receptor-Associated Factor 6 Signaling Following Traumatic Brain Injury. Front Mol Neurosci 2021, 14, 629910, doi: 10.3389/fnmol.2021.629910.

29. Lu, Y.; Cao, D.L.; Ma, L.J.; Gao, Y.J. TRAF6 Contributes to CFA-Induced Spinal Microglial Activation and Chronic Inflammatory Pain in Mice. Cell Mol Neurobiol 2022, 42, 1543–1555, doi: 10.1007/s10571-021-01045-y.

30. Masperone, L.; Codrich, M.; Persichetti, F.; Gustincich, S.; Zucchelli, S.; Legname, G. The E3 Ubiquitin Ligase TRAF6 Interacts with the Cellular Prion Protein and Modulates Its Solubility and Recruitment to Cytoplasmic p62/SQSTM1-Positive Aggresome-Like Structures. Mol Neurobiol 2022, 59, 1577–1588, doi: 10.1007/s12035-021-02666-6.

31. Ertem FB, Guven O, Buyukdag C, et al. Protocol for structure determination of SARS-CoV-2 main protease at near-physiological-temperature by serial femtosecond crystallography. STAR Protoc. 2022;3(1):101158. Published 2022 Jan 24. doi:10.1016/j.xpro.2022.101158

32. Garman, E.F.; Owen, R.L. Cryocooling and radiation damage in macromolecular crystallography. Acta Crystallogr D Biol Crystallogr 2006, 62, 32–47, doi:10.1107/S0907444905034207.

33. Atalay, N.; Akcan, E.K.; Gul, M.; Ayan, E.; Destan, E.; Ertem, F.B.; Tokay, N.; Çakilkaya, B.; Nergiz, Z.; Karakadioğlu, G.; et al. Cryogenic X-ray crystallographic studies of biomacromolecules at Turkish Light Source “Turkish DeLight”. Turkish Journal of Biology 2022, 47:1–13 doi:10.55730/1300-0152.2637

34. Rigaku. CrysAlisPro Software System, Version 1.171.42.35a. 2021, Rigaku Oxford Diffraction, https://www.rigaku.com.

35. McCoy, A.J.; Grosse-Kunstleve, R.W.; Adams, P.D.; Winn, M.D.; Storoni, L.C.; Read, R.J. Phaser crystallographic software. J Appl Crystallogr 2007, 40, 658–674, doi:10.1107/S0021889807021206.

36. Adams, P.D.; Afonine, P.V.; Bunkoczi, G.; Chen, V.B.; Davis, I.W.; Echols, N.; Headd, J.J.; Hung, L.W.; Kapral, G.J.; Grosse-Kunstleve, R.W.; et al. PHENIX: a comprehensive Python-based system for macromolecular structure solution. Acta Crystallogr D Biol Crystallogr 2010, 66, 213–221, doi:10.1107/S0907444909052925.

37. Yin, Q.; Lin, S.C.; Lamothe, B.; Lu, M.; Lo, Y.C.; Hura, G.; Zheng, L.; Rich, R.L.; Campos, A.D.; Myszka, D.G.; et al. E2 interaction and dimerization in the crystal structure of TRAF6. Nat Struct Mol Biol 2009, 16, 658–666, doi:10.1038/nsmb.1605.

38. Emsley, P.; Cowtan, K. Coot: model-building tools for molecular graphics. Acta Crystallogr D Biol Crystallogr 2004, 60, 2126–2132, doi:10.1107/S0907444904019158.

39. The PyMOL Molecular Graphics System, Version 2.5.2, Schrödinger, LLC.

40. Gul, M.; Ayan, E.; Destan, E.; Johnson, J.A.; Shafiei, A.; Kepceoglu, A.; Yilmaz, M.; Ertem, F.B.; Yapici, I.; Tosun, B.; et al. Rapid and High Resolution Ambient Temperature Structure Determination at Turkish Light Source. bioRxiv 2022, 2022.10.12.511637, doi:10.1101/2022.10.12.511637.

41. Dou, Y.; Tian, X.; Zhang, J.; Wang, Z.; Chen, G. Roles of TRAF6 in Central Nervous System. Curr Neuropharmacol 2018, 16, 1306–1313, doi:10.2174/1570159X16666180412094655.

42. Min, Y.; Kim, M.J.; Lee, S.; Chun, E.; Lee, K.Y. Inhibition of TRAF6 ubiquitin-ligase activity by PRDX1 leads to inhibition of NFKB activation and autophagy activation. Autophagy 2018, 14, 1347–1358, doi:10.1080/15548627.2018.1474995.

43. Lin Y, Bai L, Chen W, Xu S. The NF-kappaB activation pathways, emerging molecular targets for cancer prevention and therapy. Expert Opin Ther Targets 2010, 14, 45–55, doi: 10.1517/14728220903431069.

44. Cassandri, M.; Smirnov, A.; Novelli, F.; Pitolli, C.; Agostini, M.; Malewicz, M.; Melino, G.; Raschella, G. Zinc-finger proteins in health and disease. Cell Death Discov 2017, 3, 17071, doi:10.1038/cddiscovery.2017.71.

45. Krishna, S.S.; Majumdar, I.; Grishin, N.V. Structural classification of zinc fingers: survey and summary. Nucleic Acids Res 2003, 31, 532–550, doi:10.1093/nar/gkg161.

46. Cai, C.; Tang, Y.-D.; Zhai, J.; Zheng, C. The RING finger protein family in health and disease. Signal Transduction and Targeted Therapy 2022, 7, doi:10.1038/s41392-022-01152-2.

47. Otsuka, M.; Fujita, M.; Aoki, T.; Ishii, S.; Sugiura, Y.; Yamamoto, T.; Inoue J. Novel zinc chelators with dual activity in the inhibition of the kappa B site-binding proteins HIV-EP1 and NF-kappa B. J Med Chem 1995, 38, 3264–3270, doi: 10.1021/jm00017a011.

48. Otsuka, M.; Fujita, M.; Sugiura, Y.; Yamamoto, T.; Inoue, J.; Maekawa, T.; Ishii, S. Synthetic inhibitors of regulatory proteins involved in the signaling pathway of the replication of human immunodeficiency virus 1. Bioorg Med Chem 1997, 5, 205–215, doi: 10.1016/s0968-0896(96)00203-9.

49. Fujita, M.; Otsuka, M.; Sugiura, Y. Metal-chelating inhibitors of a zinc finger protein HIV-EP1. Remarkable potentiation of inhibitory activity by introduction of SH groups. J Med Chem 1996, 39, 503–507, doi: 10.1021/jm950831t.

50. Ejima, T.; Hirota, M.; Mizukami, T.; Otsuka, M.; Fujita, M. An anti-HIV-1 compound that increases steady-state expression of apoplipoprotein B mRNA-editing enzyme-catalytic polypeptide-like 3G. Int J Mol Med 2011, 28, 613–616, doi: 10.3892/ijmm.2011.737.

51. Koga, R.; Radwan, M.O.; Ejima, T.; Kanemaru, Y.; Tateishi, H.; Ali, T.F.S.; Ciftci, H.I.; Shibata, Y.; Taguchi, Y.; Inoue, J.I.; Otsuka, M.; Fujita, M. A Dithiol Compound Binds to the Zinc Finger Protein TRAF6 and Suppresses Its Ubiquitination. ChemMedChem 2017, 12, 1935–1941, doi: 10.1002/cmdc.201700399.

52. Tanaka, A.; Radwan, M.O.; Hamasaki, A.; Ejima, A.; Obata, E.; Koga, R.; Tateishi, H.; Okamoto, Y.; Fujita, M.; Nakao, M.; Umezawa, K.; Tamanoi, F.; Otsuka, M. A novel inhibitor of farnesyltransferase with a zinc site recognition moiety and a farnesyl group. Bioorg Med Chem Lett 2017, 27, 3862–3866, doi: 10.1016/j.bmcl.2017.06.047.

53. Radwan, M.O.; Koga, R.; Hida, T.; Ejima, T.; Kanemaru, Y.; Tateishi, H.; Okamoto, Y.; Inoue, J.I.; Fujita, M.; Otsuka, M. Minimum structural requirements for inhibitors of the zinc finger protein TRAF6. Bioorg Med Chem Lett 2019, 29, 2162–2167, doi: 10.1016/j.bmcl.2019.06.050.

54. Tateishi, H.; Tateishi, M.; Radwan, M.O.; Masunaga, T.; Kawatashiro, K.; Oba, Y.; Oyama, M.; Inoue-Kitahashi, N.; Fujita, M.; Okamoto, Y.; Otsuka, M. A new inhibitor of ADAM17 composed of a zinc-binding dithiol moiety and a specificity pocket-binding appendage. Chem Pharm Bull 2021, 69, 1123–1130, doi: 10.1248/cpb.c21-00701.

