## Supplementary figures and images for "Structural Characterization of TRAF6 N-terminal for Therapeutic Uses"

### https://drive.google.com/drive/u/0/folders/1GejcUMXvZ9ryu4zG1ClzIVw0CU0_s54A

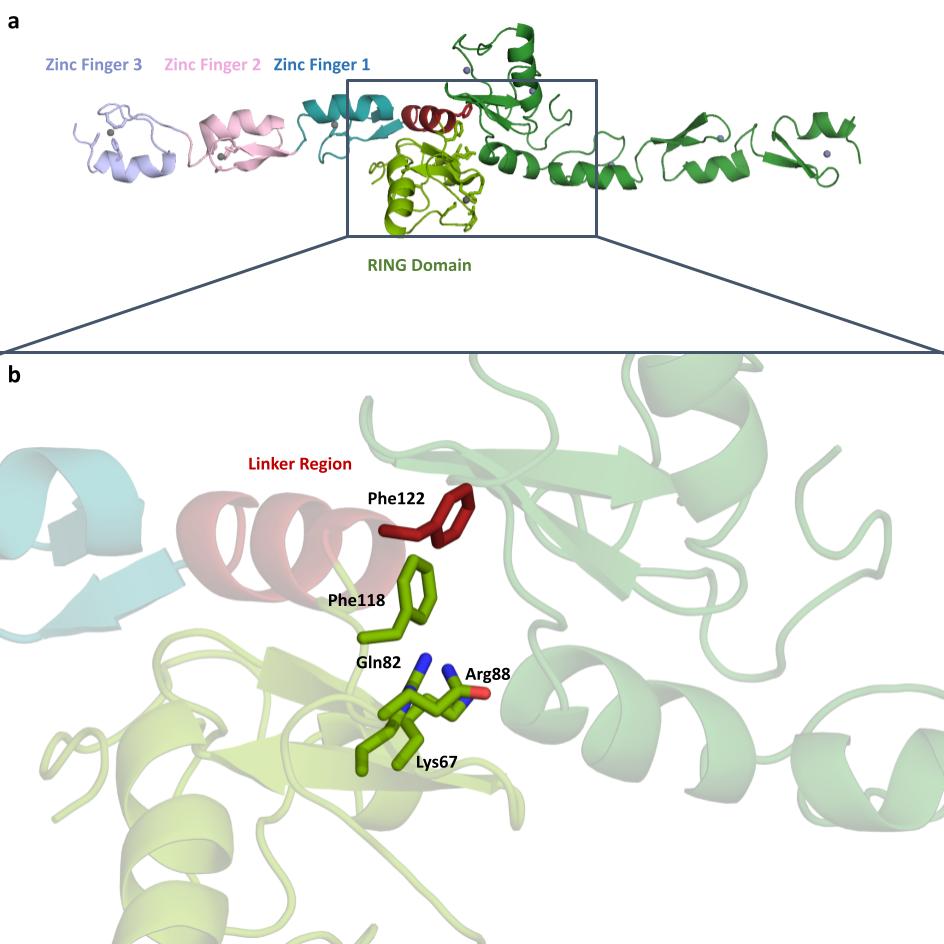
